# CSL-Tox: An open-source analytical framework for the comparison of short-term and long-term toxicity end points and exploring the opportunities for decreasing in-vivo studies conducted for drug development programs

**DOI:** 10.1101/2022.08.06.503048

**Authors:** Doha Naga, Smaragda Dimitrakopoulou, Sonia Roberts, Elisabeth Husar, Susanne Mohr, Helen Booler, Eunice Musvasva

## Abstract

In-vivo toxicity assessment is an important step prior to clinical development and is still the main source of data for overall risk assessment of a new molecular entity (NCE). All in-vivo studies are performed according to regulatory requirements and many efforts have been exerted to minimize these studies in accordance with the (Replacement, Reduction and Refinement) 3Rs principle. Many aspects of in-vivo toxicology packages can be optimized to reduce animal use, including the number of studies performed as well as study durations, which is the main focus of this analysis. We performed a statistical comparison of adverse findings observed in 116 short-term versus 78 long-term studies in order to explore the possibility of using only short-term studies as a prediction tool for the longer-term effects. Annotation of treatment related findings was one of the challenges faced during this work. A specific focus was therefore put on the summary and conclusion sections of the reports since they contain expert assessments on whether the findings were considered adverse or were attributed to other reasons. Our analysis showed a general good concordance between short-term and long-term toxicity findings for large molecules and the majority of small molecules. Less concordance was seen for certain “target organ systems findings’. While this work supports the minimization of in-vivo study durations, a larger-scale effort would be needed to provide more evidence. We therefore present the steps performed in this study as an open-source R workflow (CSL-Tox) and we provide the dataset used in the work to allow researchers to reproduce such analysis and to promote large-scale application of this study.

## I. Introduction

Adverse drug reactions (ADRs) represent one of the major causes of drug attrition [1–4] and have previously caused several market withdrawals [5–8]. With the purpose of identification and elimination of such hazardous adverse events in the development of new medicines, testing starts prior to studies in animals; safety pharmacology (SP) and *in vitro* studies are followed by *in vivo* toxicity studies prior to entry in humans [9, 10]. *In vivo* toxicity studies are pivotal to support clinical studies at all stages up to registration. The required non-clinical safety package varies among molecule classes, but the principles are similar [11–13]. One of the main purposes of *in vivo* toxicity studies is identification of the target organ specific toxicities, providing a safety margin and exploring the reversibility of the adverse events observed. This information is used to estimate an initial safe starting dose and dose range for human trials and to identify parameters for clinical monitoring of potential adverse effects. Investigation of several parameters that cannot be obtained in clinical trials, such as macroscopic and microscopic pathological findings, help in the identification of toxicity. They also help in the determination of the highest dose that does not produce a significant increase in adverse events and does not raise any safety concerns, defined as the No Observed Adverse Effect Level (NOAEL) [14].

In-vivo toxicity experiments are performed in accordance with the replace, refine and reduce (3Rs) principles [15, 16]. Sparrow *et al* reviewed the study designs of cross-company toxicity studies and highlighted the factors affecting the number of animals used, such as the general design of the toxicology program and the use of control groups [17]. They proposed case-specific adjustments in study designs that allow for minimization of animal use. The use of virtual control groups was also proposed [18]. Other efforts questioned the utility of two species in a toxicity study, proposing the use of only one species in long-term toxicity studies. All these efforts contribute to the minimization of the number of animals used in toxicity studies [19, 20].

At Roche *in vivo* experiments are conducted in compliance with the highest international standards for animal welfare. To reduce the number of non-rodent animals, studies are often combined, the use of recovery groups is avoided whenever possible, control animals are not used in dose range finding studies unless there is a clear scientific need and non-human primates are re-used for pharmacokinetic and dose range finding studies.

Reduction of the number of studies performed is another aspect serving the 3Rs principle and is the focus of this work. Toxicity can become manifest either after a short time or only after repeated exposure to the drug throughout longer durations. The recommended duration of repeated-dose toxicity studies is usually related to the duration, therapeutic indication and scope of the proposed clinical trial. Fig.1 represents an overview of the different types of toxicity studies performed during the drug development process and the corresponding durations, which are dependent on the clinical program. Generally, repeated-dose toxicity studies for a minimum duration of 2 weeks would generally support any clinical development trial up to 2 weeks in duration. Clinical trials of longer duration should be supported by repeated-dose toxicity studies of at least equivalent duration. 6-month rodent and 9-month non-rodent studies generally support dosing for longer than 6 months in clinical trials (see ICH M3 guideline) [11]. Toxicity studies can be classified into short-term studies e.g. acute toxicity, short-term studies, sub-chronic studies that generally last for 1 to 3 months and long-term studies e.g. chronic studies that may last up to 12 months. Generally, long-term studies can indicate the progression of findings that were either not observed in shorter studies or were of low incidence, or less severe and therefore not considered adverse. However, this might not always be the case and sometimes no new adverse events are observed upon increasing the study duration. The likelihood of identifying new adverse events with longer durations is therefore explored in this study. This might provide an opportunity to remove some of the long-term studies, therefore reducing the number of studies performed and consequently the overall number of animals in in-vivo studies. Upon the analysis of 59 sub-chronic and chronic studies, the authors concluded that most toxicities were identified in studies up to 3 months duration and that compound termination in the clinical phase due to new findings in chronic toxicity studies were rare (about 10%) [21]. Other publications comparing toxicity end points of short and long term studies, were limited to large molecules [22, 23].

**Figure 1.**
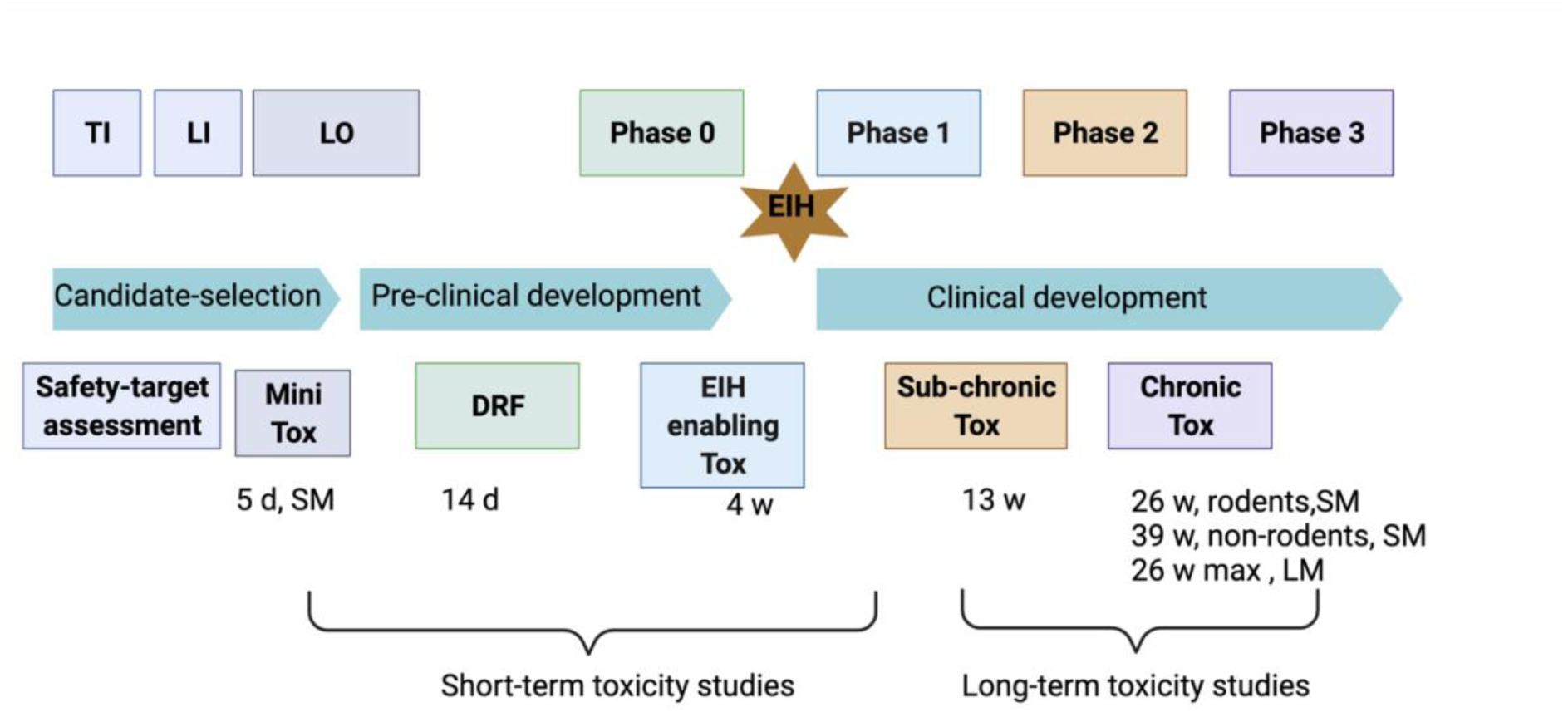
Toxicological testing of new molecular entities in the different stages of drug discovery and development.

This paper presents an open-source R workflow, termed CSL-Tox, which facilitates the comparison between acute and chronic toxicity studies, with respect to the adverse findings reported. In this study, a large cohort of *in vivo* toxicological rodent and non-rodent studies were analyzed (a total of 192 studies) with representation across both small and large molecules (25 and 18 molecules respectively). Throughout this comparison, we explore the sufficiency of short-term studies in detecting adverse findings and the corresponding NOAELs. Bayesian statistics and likelihood ratios, which are commonly used methods in non-clinical and clinical concordance studies, were adopted in this comparison [24–26]. The open-source R program is available for application to other similar datasets. This can help researchers in analyzing and comparing additional short-term and long-term studies and therefore contribute to the minimization of study durations.

Finally, we discuss challenges faced during this study such as cumbersome data extraction and annotation of treatment related findings and current efforts to overcome these challenges.

## I. Methods

### Workflow

An overall view of the work is given in Figure 1. The work can be summarized into three main steps: (1) data collection and studies overview (modality, species, duration); (2) data refinement (including the high-level categorization of the findings, converting the finding terms into the controlled terminology and flagging the treatment related findings); (3) Data analysis of the results (calculation of the overall adversity, NOAEL changes and the Bayesian contingency tables). The functions corresponding to the steps explained in this workflow and implemented in the software are provided in the “Implementation and tools” section (**Table. 3**).

### 1-Data collection and overview of the studies

In-house study report databases were searched for general toxicity studies conducted on rodent and non-rodent species. Both modalities, small and large molecules were included. No restriction was placed on the timeframe for the studies collected. However, for a molecule to be considered in the database, short or/and mid and long-term studies should have been conducted. Therefore, the main inclusion criteria of the molecules in the database were based on the study duration:

#### (a) Short-term and midterm studies

A general toxicity study with a duration of less than 20 weeks should have been conducted. We further distinguish between two classes of studies: general toxicity studies with a duration of up to a maximum of 6 weeks (short-term) and studies with a duration of more than 6 weeks and less than 20 weeks (midterm).

#### (b) Long-term studies

A general toxicity study with a duration of at least 26 weeks should have been conducted (long-term). According to the current guidelines, those studies are required to support continued clinical development and marketing of molecules, hence they are conducted at the latest stage of the drug development process. As a direct consequence, molecules that were terminated in initial stages due to adversity or other commercial reasons were excluded from the database. Furthermore, this requirement resulted in the exclusion of anticancer pharmaceuticals that were developed after 2010, when the revised ICHS9 guideline restricting the maximum duration of a nonclinical study for advanced cancer indications to 13 weeks, came into effect [13].

Information on the study and the therapeutic molecules were gathered from the study reports such as the study duration, species, modality of the molecule, route of administration, dose interval, no observed adverse effect level and adversity or treatment-related findings were registered in the database. The latter two can be described as follows:

##### No observed adverse effect level (NOAEL)

NOAEL is defined as the highest dose administered at which no adverse effects are observed [27]. The NOAEL can be a representative measure of how the toxicity, associated with a specific treatment, progresses as the duration of the study increases and was therefore considered in this work. For an overall analysis of the effect of the study duration on the NOAEL, short-term and midterm studies for each species have been grouped together and the lowest specified NOAEL has been identified per molecule. This was used as a basis to assess the influence of long-term treatment on the NOAEL. If more than one long-term study was conducted for a specific species, the lowest of the defined NOAELs was used in the comparison.

##### Adverse treatment-related findings

“Treatment related” findings are flagged by the study responsible as a finding that is directly related to the drug. “Adverse” is a term indicating “harm” to the test animal, while non-adverse indicates lack of harm. The “Summary” and “Conclusion” section of the study reports were the main source for the extraction of toxic effects that were considered treatment-related. Detailed information on treatment-related findings (categories specified below) recorded in the toxicity study was collected. Specifically, the dose at which the effect appeared, and most importantly whether it was considered adverse by the toxicologist in charge. The level of severity and reversibility and any extra information related to the effect or its cause (if stated, e.g. whether it was considered secondary to stress) were also recorded.

The findings were grouped into 9 broad categories. The categories, as well as examples of effects/events belonging to each of the categories are shown in Table 1. The full details on the events included in each category (except for microscopic pathology) is provided in the supplementary material (**Table S1-A, B, C, D, E**). The detailed microscopic pathology events are provided in the supplementary material (**Table S2**). For the effects belonging to the categories “organ weights”, “macroscopic pathology” and “microscopic pathology” their target organs were also recorded. Information on toxicokinetics were not included as they were considered out of scope for this particular analysis.

**Table 1.** Summary of the 9 adverse events categories further aggregated into 6 high level categories. Examples of events are provided for each category. Full details on the events are provided in **Table S1** and **S2**.

| Categories | High level categories | Event examples |
| --- | --- | --- |
| Body weight | Body weight changes | increase, decrease |
| Body weight gain |  | increase, decrease |
| Macroscopic pathology | Post-mortem findings<br>(Target organ system) | decrease in size (thymus gland) |
| Microscopic pathology |  | hepatocyte hypertrophy (liver) |
| Organ weights |  | Decrease (liver) |
| Cardiovascular effects | Cardiovascular effects | Increased heart rate, QTc prolongation |
| Neurological clinical signs | Neurological clinical signs | hypoactivity, tremors |
| GIT clinical signs | GIT clinical signs | salivation, vomit |
| Other clinical signs | Other clinical signs | irregular respiration |

### 2-Data refinement (high level effect aggregation and controlled terminology)

Firstly, species were grouped into rodents (rat and mouse) and non-rodents (cynomolgus monkey, minipig, marmoset and dog). Secondly doses were normalized for each study i.e. since dosing intervals may vary between studies, doses were computed per day in order to be able to compare different studies). Finally, since many of the toxicity effects showed low prevalence, grouping of the effects was performed in order to gain a better understanding of the overall incidence of adverse events observed in long-term, midterm and short-term studies. No grouping was performed for the four categories: neurological clinical signs, other clinical signs, gastro-intestinal tract and cardiovascular effects. The findings within each category are presented in detail in Table S1. A, B, C, D respectively. On the other hand, the three categories macroscopic pathology, microscopic pathology and organ weights were grouped into “post-mortem findings” and, the two categories body weight and body weight gain, were grouped into “body weight changes” as follows:

#### Post-mortem findings (Target-organ system)

The “macroscopic pathology”, “microscopic pathology” and “organ weights” categories were merged into one and the effects were associated with the target organ system affected. For example, all the postmortem effects belonging to the mentioned categories and affecting organs in the Endocrine system were grouped under the high-level effect “Endocrine system”. The target systems grouping was performed according to Table S2, where it is described in detail which organs or tissues constitute each target system. The macroscopic pathological findings are described in Table S1-E and the microscopic findings are described separately in Table S2 due to the considerable amount of information.

#### Body weight changes

“Body weight” and “body weight gain” categories were combined into one category named “body weight changes” since both reflect the influence of the therapeutic molecule on the weight of the animals.

A summary of the presence/absence of the findings within the described categories was included in the dataset.

#### Overall adversity measure

Based on the previous categories, the “overall adversity” of each compound was registered in the database, where it was labelled as “No” if zero adverse event was observed and “Yes” if at least one adverse event was observed.

### 3-Data analysis and statistical methods

The statistical methods implemented in this study and used for the analysis of the adverse events were adopted from the work of Clark *et al.*^23^. We used Bayesian statistics with a 2 by 2 contingency table to measure the concordance between observations made in the initial short and midterm general toxicity studies and the long-term toxicity studies. We treat the observations made in the early toxicity studies as a diagnostic test for observations made in the long-term toxicity studies and use the statistical methods developed for the evaluation of the efficacy of these diagnostic tests. The contingency tables were calculated after the high level grouping of adverse events (explained previously) was performed.

The values in the contingency table, which represent number of molecules in each of the four categories for a specific high-level effect are explained in **Fig.3** and were generated as follows:

(a) True positives: Count of molecules for which the high-level effect was observed in the long-term study as well as in the short-term or/and mid-term study.

(b) False positives: Count of molecules for which the high-level effect was observed in the short-term or/and mid-term study but not in the long-term study

(c) True negatives: Count of molecules for which the high level effect was not observed in any study (neither short/mid nor long term)

(d) False negatives: Count of molecules for which the high-level effect was not recorded in the short-term or/and mid-term study but was recorded in the long-term study

#### Likelihood ratios

Likelihood ratios were used to determine the statistical connection between the toxicity observations made in the initial toxicity studies and the long-term study. Likelihood ratios were considered the appropriate metric since they combine the knowledge of both the sensitivity and the specificity of the model^21^. Also, they have the advantage that they are independent from the prevalence, allowing for the comparison of high level effects with different prevalence in the dataset.

The positive likelihood ratio (LR+) is defined as sensitivity /(1-specificity) whereas the inverse Negative Likelihood (iLR-) is defined as specificity/(1-sensitivity). Likelihood ratios were calculated according to equation 3 and 4 respectively. The p-values for the relationships in the 2 by 2 contingency tables were computed using Fisher’s exact test and the interpretation of likelihood ratios were adopted from Chien and Khan’s work and is listed in Table.2 ^16^. Only statistically significant likelihood ratios (p-value < 0.05) were considered in the analysis and a cut-off of 5 was drawn to indicate a “high” positive or inverse negative likelihood ratio.

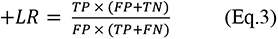

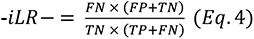

#### Frequency or percentage of false positives and false negatives

Miss-predictions of findings suggested by the short-term studies were estimated by the percentage/frequency of false positives (the events that were observed in short-term studies and not observed in long-term studies, calculated by Eq.5) and the percentage or frequency of false negative (the events that were observed in long-term studies and missed by the short-term studies, calculated by Eq.6).

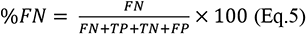

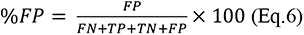

### Implementation and tools

All the data manipulation and statistical analysis was implemented in R studio (version 3.5.1) [28]. Figures 1 and 2 were created using www.biorender.com and Package ggplot (version 3.3.2) was used for plotting figures 3-5. The CSL-Tox code and tutorial are available at the Github repository: https://github.com/Roche/CSL-Tox.

**Figure 2.**
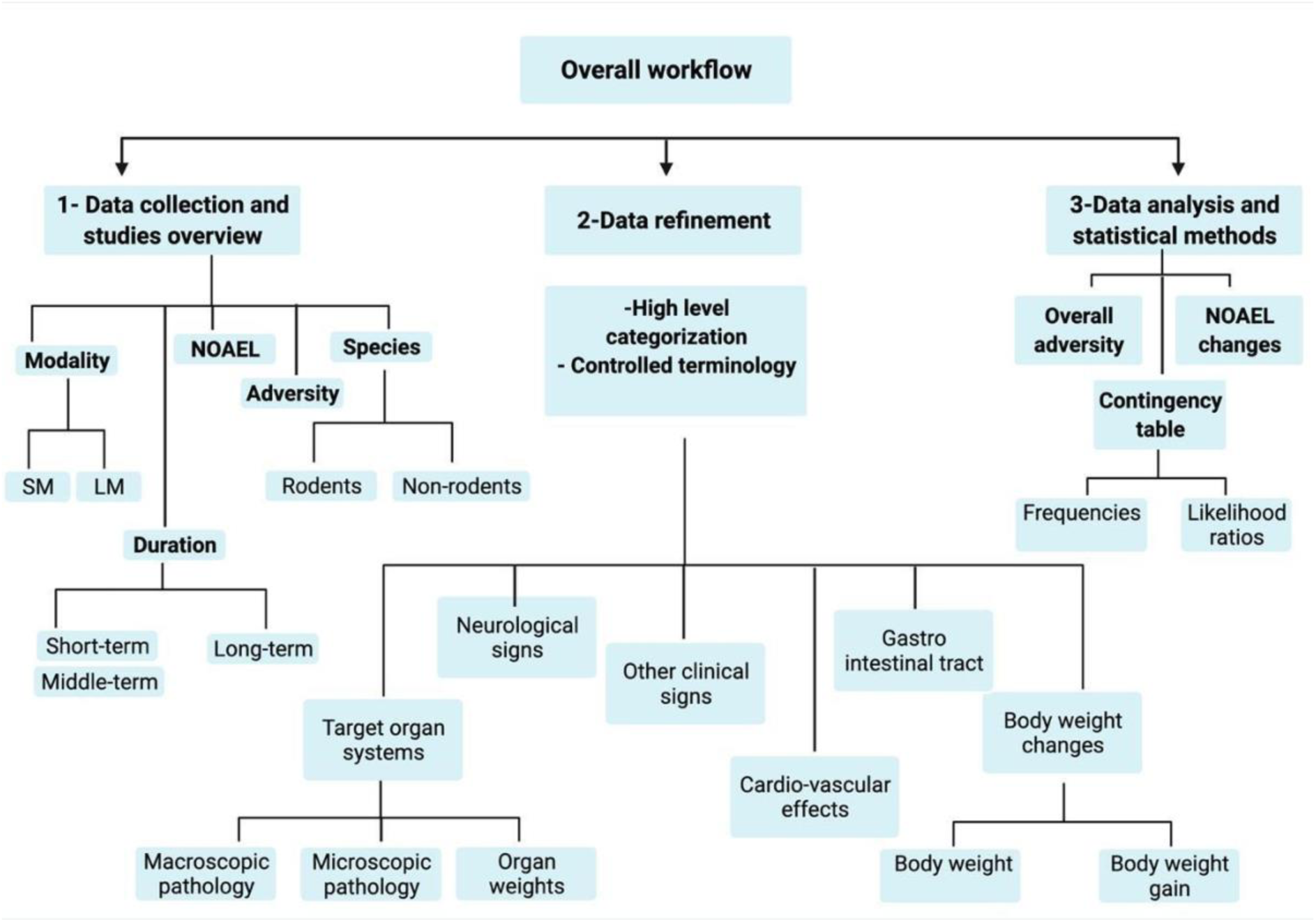
An overview of the steps performed in this work and implemented in the “CSL-Tox ” workflow.

**Figure 3.**
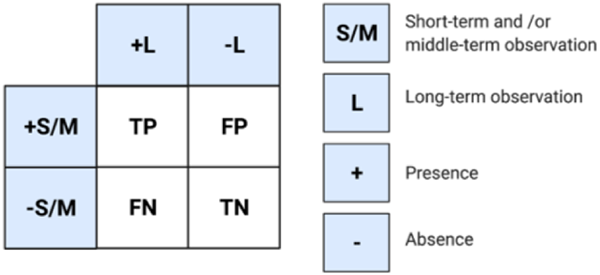
Contingency table used in the statistical analysis of the findings.

## II. Results

### Overview of the dataset

A representation of the compiled dataset is given in **Fig.4** with respect to the therapeutic areas covered by the compounds, the study durations, the type and number of species for each study and the overall distribution of adverse events for rodents and non-rodents.

**Figure 4.**
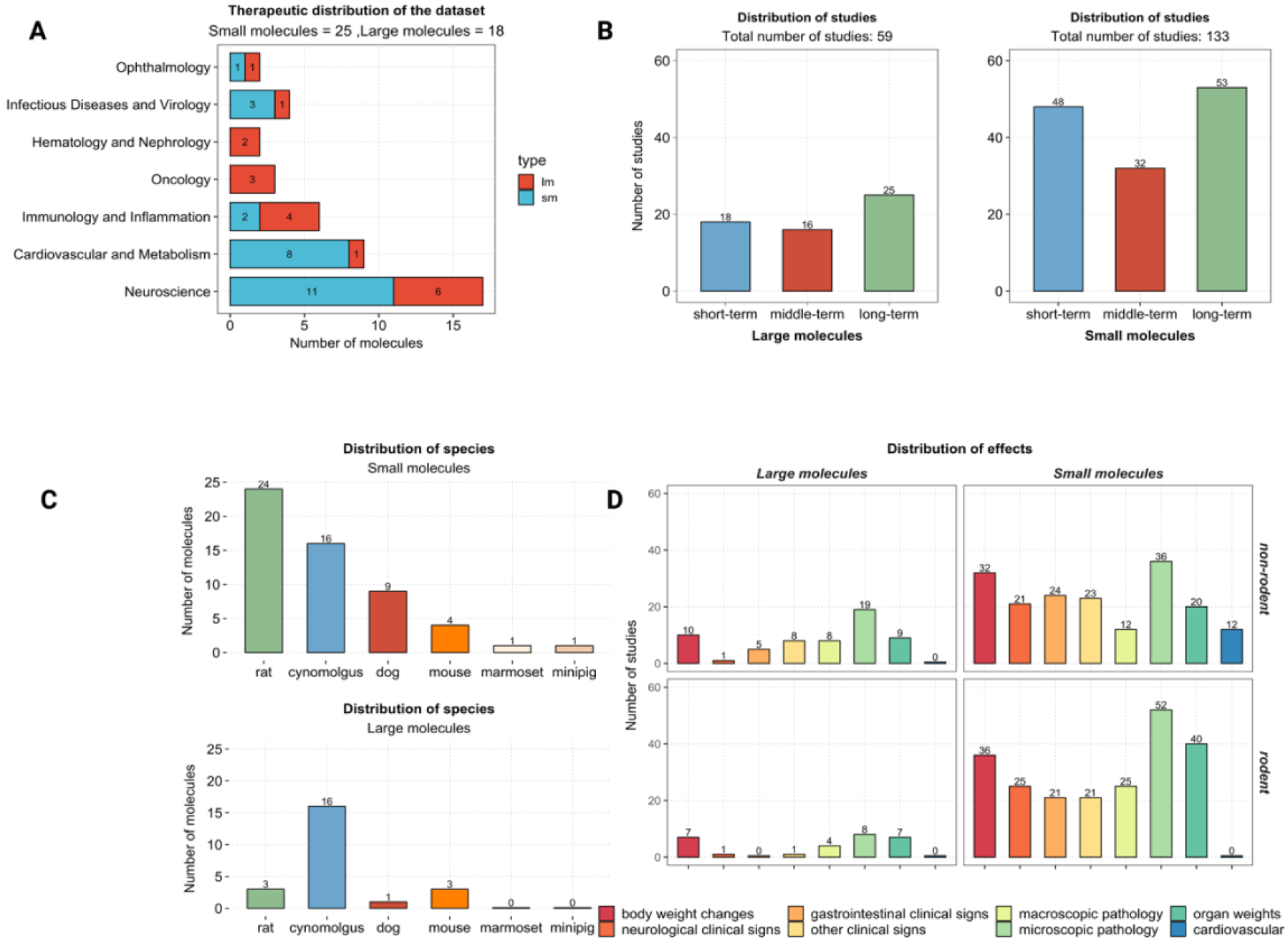
An overall illustration of the compiled dataset with respect to therapeutic areas covered, study durations, species tested and adverse events observed. (A) A bar plot showing therapeutic areas covered by the dataset, the number of molecules belonging to each therapeutic area is indicated on the corresponding bar. Each bar is colored according to the type of molecule. (B) A bar plot showing the number of studies with respect to the duration (short, mid and long term). The number of studies present in each duration category is indicated on the top of the corresponding bars. (C) A bar plot showing the number of compounds tested per species (for rodents and non-rodents). (D) A bar plot showing the number of studies for which adverse events were registered in rodent and non-rodent species (number of studies is indicated on the top of each bar) . The bars are colored according to the adverse event categories.

A total of 43 molecules met the inclusion criteria (including 25 small molecules and 18 large molecules), representing a wide range of therapeutic areas as shown in **Fig.4A**, with “Neurosciences” being the most represented area. The full dataset is provided in the supplementary material (**Table. S3, Table.S4**).

The distribution of the studies across the different specified duration classes is shown in **Fig. 4B**. For the large molecules, a total number of 61 general toxicity studies are included in the dataset (25 long-term studies). As for small molecules, a total number of 133 general toxicity studies are included in the dataset (53 long-term studies). A short-term and a midterm study were not always conducted as part of a new molecule’s toxicity testing, with the short-term study being preferred to the midterm study.

In **Fig.4C**, the bar plots show the number and type of species used in the toxicity studies for both small and large molecules, while the pie charts indicate the number of species used according to the current ICH guidelines.

For most of the small molecules (20/25), general toxicity studies were conducted in two species, one rodent and one non-rodent, as recommended by the current ICH M3(R2) guideline and indicated in the pie chart of Fig 3C. For one-fifth of the small molecules, a further third species was used to test the molecule’s toxicity. As seen in the figure, the rat was the predominant rodent species (24/25) while cynomolgus (16/25) followed by the dog (9/25) were the main non-rodent species.

With regards to large molecules, the current guideline for large molecules, ICH S6(R1), recommends the testing of novel medicinal products in pharmacologically relevant species only; if possible, a rodent and non-rodent. Frequently the only such species is the cynomolgus monkey, due to the higher sequence homology between the human and cynomolgus monkey proteome. Therefore, general toxicity studies were conducted in only one species for the majority of large molecules (13/18 molecules), whereas a second relevant species was used in only 5 out of the 18 molecules. Cynomolgus monkey was the predominant selected non-rodent species (16/18). One molecule was only tested in the dog and one molecule was only tested in the mouse. The rat was the selected rodent species for 3 large molecules, whereas the mouse was selected for the remaining 2 molecules.

Fig. 4D shows the distribution of all the findings recorded within the studies in the specified categories. The frequency of findings observed for large molecules was lower than that for small molecules, which can be attributed to the higher selectivity of large molecules. “Microscopic pathology” followed by “organ weights” and “body weight changes” were the categories with the most frequently observed effects in both small molecules and large molecules.

### Overall adversity

As explained in the “Methods” section, an overall adversity measure was assigned to gain an overall understanding on the concordance of findings between the short/mid-term studies and the long-term studies. The overall adversity observations are displayed in Table 4 for large molecules, where a concordance between short/mid-term and long-term studies is observed for the majority of the molecules (16 out of 18). 14 out of the 16 molecules that showed concordance did not show any adverse events in neither the short/mid nor the long-term studies. Three out of 18 molecules showed adverse events in both the short/mid and the long-term studies. For one out of the 18 large molecules (Compound-O) an antibiotic resistant infection, resulting in a skin lesion, was observed in one animal in the long-term studies and not in the short-term study. The short-term study is not considered fully unpredictive in this case since it was unclear if the skin infection was compound related. Large molecules therefore showed an overall high concordance in the findings between short and long-term studies.

**Table 2.** Interpretation of likelihood ratios

| +LR and -iLR | Significance |
| --- | --- |
| >10 | Large and often conclusive shifts in probability |
| 5-10 | Moderate shifts in probability |
| 2-5 | Small, but sometimes important, shifts in probability |
| 1-2 | Alters probability to a small, and rarely important, degree |

**Table 3.** Key functions available in the workflow.

| Category | Functions | Description |
| --- | --- | --- |
| Data exploration | TherapeuticAreas<br>DifferentSpecies<br>StudiesPerFindingCat | Plots an overview of the therapeutic areas, species and findings available in the dataset |
| Data refinement | ReadingData | Reads data from a .txt file and put it in the required format |
|  | ChangeData | Applies controlled terminology (Table S1, S2) to the 9 categories of findings and groups them into the 6 high level categories explained in Table.1 |
| Data analysis | AdversitySummary | Calculates the overall adversity per modality (small molecules and large molecules). |
|  | NOAELChange | Calculates NOAEL increase, decrease or same from short and middle to long-term studies. The lowest NOAEL doses are compared in case of multiple studies. |
|  | LikelihoodRatio | Calculates contingency table, +LR, -iLR and corresponding Fisher test p-values per species (rodents or non-rodents) |

**Table 4.** A comparison between the occurrence of adverse events in large molecules for the short and mid-term studies versus long-term studies per compound.

| Large molecule | Short or mid-term study | Long-term study |
| --- | --- | --- |
| Compound- A to Compound-N | No adverse events observed |  |
| **Compound-O** | No adverse events observed | Adverse events observed |
| Compound-P to Compound-R | Adverse events observed |  |

For small molecules, the short/mid-term toxicity studies were able to predict the adverse events in long-term studies for the majority of the molecules (18 out of 25) as seen in Table 5. A difference in findings between the short/mid- and long-term studies was seen for 6 out of 25 small molecules, only 2 of which (Compound-6 and Compound-7) where adverse events were captured only in the long-term studies. For Compound-6 [4w, 12w and 39 w], similar findings were observed in the 4w and 12 study (thyroid and parathyroid findings in 4 w study were reversible and appeared in 12 w study). However, in the 39w study other adverse events were observed, such as decreased prostate weights. Animals were generally younger in age at study start in long-term studies and effects on the prostate were only observed as animals matured. Short term studies in older animals may have missed this. For Compound-7 [4w, 26w] axonal degeneration was observed in the long-term study and was minimal. Other findings such as food consumption changes were predicted by the short-term studies.

**Table 5.** A comparison between the occurrence of adverse events in small molecules for the short and mid-term studies versus long-term studies per compound.

|  | Short or mid-term study | Long-term study |
| --- | --- | --- |
| Compound-1 to Compound-5 | No adverse events observed |  |
| **Compound-6 and Compound-7** | No adverse events observed | Adverse events observed |
| Compound-8 to Compound 11 | Adverse events observed | No adverse events observed |
| Compound 12 to Compound 25 | Adverse events observed |  |

### Progression of the “No observed adverse effect levels” (NOAEL) with study duration

As previously mentioned, the determination of a NOAEL is a very important objective of preclinical toxicity studies^2^. Since toxicity may progress with longer duration of exposure (with respect to incidence and severity), we were interested in understanding NOAEL changes from short-term to long-term studies. As seen in **Table. 6**, for small molecules, a NOAEL was identified for non-rodent studies in 23 molecules; where 8 decreased, 12 stayed the same and 3 increased from short-term to long-term studies. In rodent studies a NOAEL was identified for 22 molecules; where 10 decreased, 7 stayed the same, 5 increased.

**Table 6.**
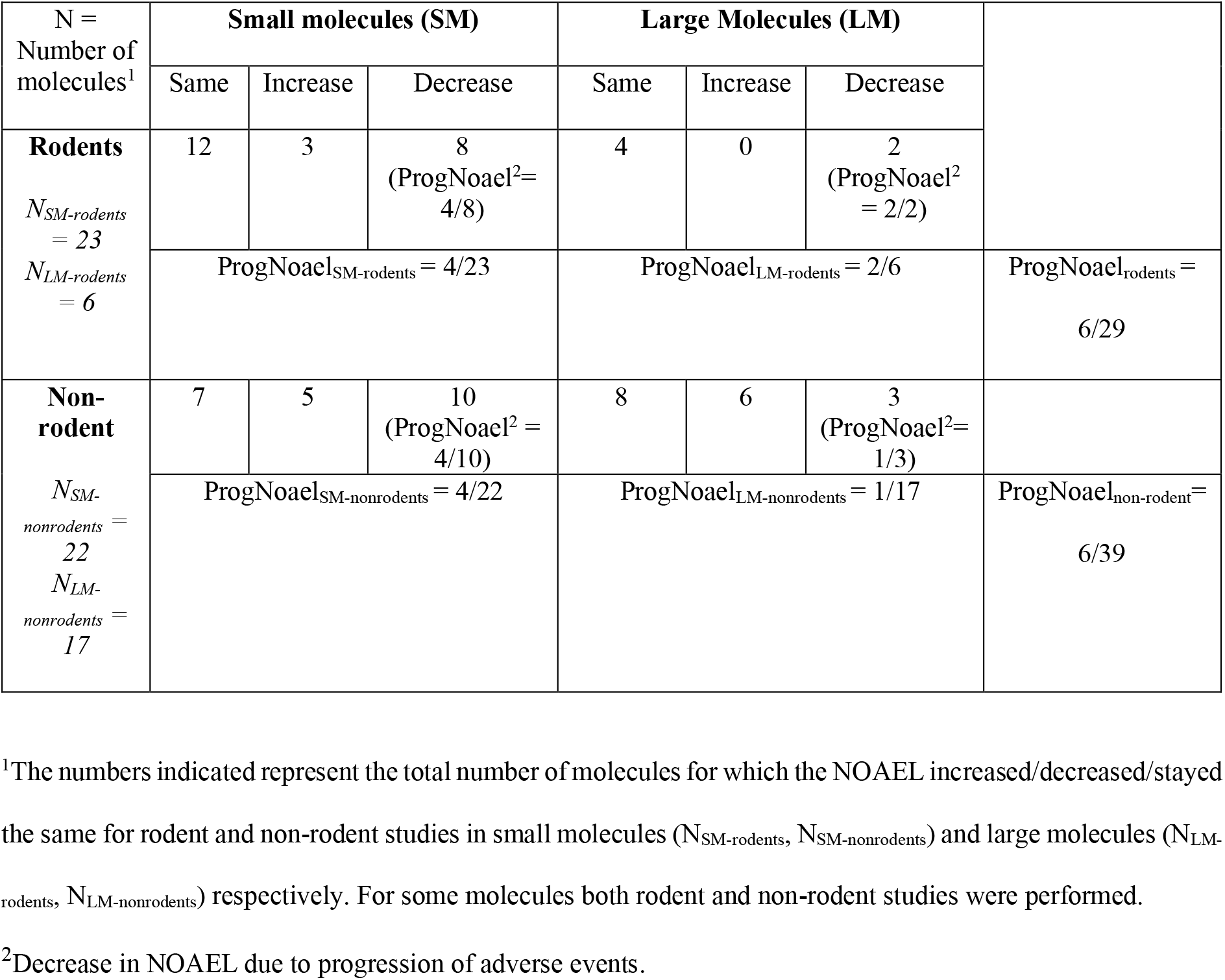
NOAEL changes for small and large molecules in rodent and non-rodent studies with a focus on NOAEL decrease attributed to progression of adverse events.

For large molecules, a NOAEL was identified in for 6 large molecules and decreased for 2 molecules, and it stayed the same for 4 molecules from short-term to long-term studies.

In rodent studies a NOAEL was identified for 17 molecules and the NOAEL stayed the same or increased for the majority of the molecules (stayed the same for 8 molecules, increased for 6 molecules) and decreased for 3 molecules.

For both, small and large molecules, the NOAEL increase was attributed to higher doses tested in longer-term studies. The NOAEL decreased in the longer-term studies, due to progression of findings for 4 out of 23 small molecules and 2 out of 6 large molecules in rodent studies. In non-rodent studies, the NOAEL decrease was attributed to progression of adverse events for only 1 out 17 large molecules and 4 out of 22 small molecules. Otherwise, the NOAEL decrease was due to lower doses tested in the longer duration studies.

In conclusion, the overall NOAEL decrease due to progression of adverse events was seen for only 20% (6 out of 29) and 15.3% (6 out of 39) of the molecules in rodent and non-rodent studies respectively. It is worth to say that for some compounds these findings had already been observed in the shorter duration studies but were not considered adverse due to low incidence or severity.

As explained in the methods section, the NOAEL change algorithm compares the NOAEL raw values of the short and middle vs long-term studies. The results included in Table. 4, Table. 5 Table. 6 are the NOAEL changes after the toxicologists’ revisions of the algorithm outcome and interpretation of the study reports. Few discrepancies were observed between the algorithm outcome and the toxicologists’ interpretations in the NOAEL changes (provided in the supplementary material (**Table. S5**)). For example, algorithm expected an increase in the NOAEL Compound K in rodents and non-rodents while the toxicologist interpreted the NOAEL as staying the same. The “errors” identified by the toxicologists in the NOAEL algorithm changes were attributed to:

(1) Changes in some studies were irrelevant to human, therefore the toxicologist disregarded these studies in the NOAEL change estimation.

(2) Changes were due to differences in the study design and not due to better/worse tolerability in long term studies.

(3) Difference in the definition of adversity of a molecule from one toxicologist to another (due to better acquaintance with the molecule for example). Since adversity is driven by findings that have functional consequences on organs, increased knowledge of presence or absence of functional consequences may also change adversity definitions from a pathologist’s perspective.

### Positive and negative likelihood ratios

Positive and negative likelihood ratios of both adverse and non-adverse findings are given in Table. 7 and Table. 8 providing an overview on the ability of short-term studies to predict the appearance or non-appearance of adverse findings (or non-adverse findings) in the long-term studies. For example, a high positive likelihood ratio indicates that there is a high probability to observe adverse events in long-term studies, given that they appear in the short-term studies. A high inverse negative likelihood ratio indicates that there is a high probability of not observing the adverse events in long-term studies if they were not observed in the short-term studies. Therefore, the high positive or inverse likelihood ratios demonstrate the concordance of findings between short-term and long-term studies. A high +LR (>5) was seen for the majority of findings for non-rodents (Table. 7a) except for weight changes, other clinical signs and lymphoid tissue findings which showed a lower +LR (∼4). More importantly, very high likelihood ratios were observed for all the adverse findings in non-rodents (Table. 7b).

**Table 7a.**
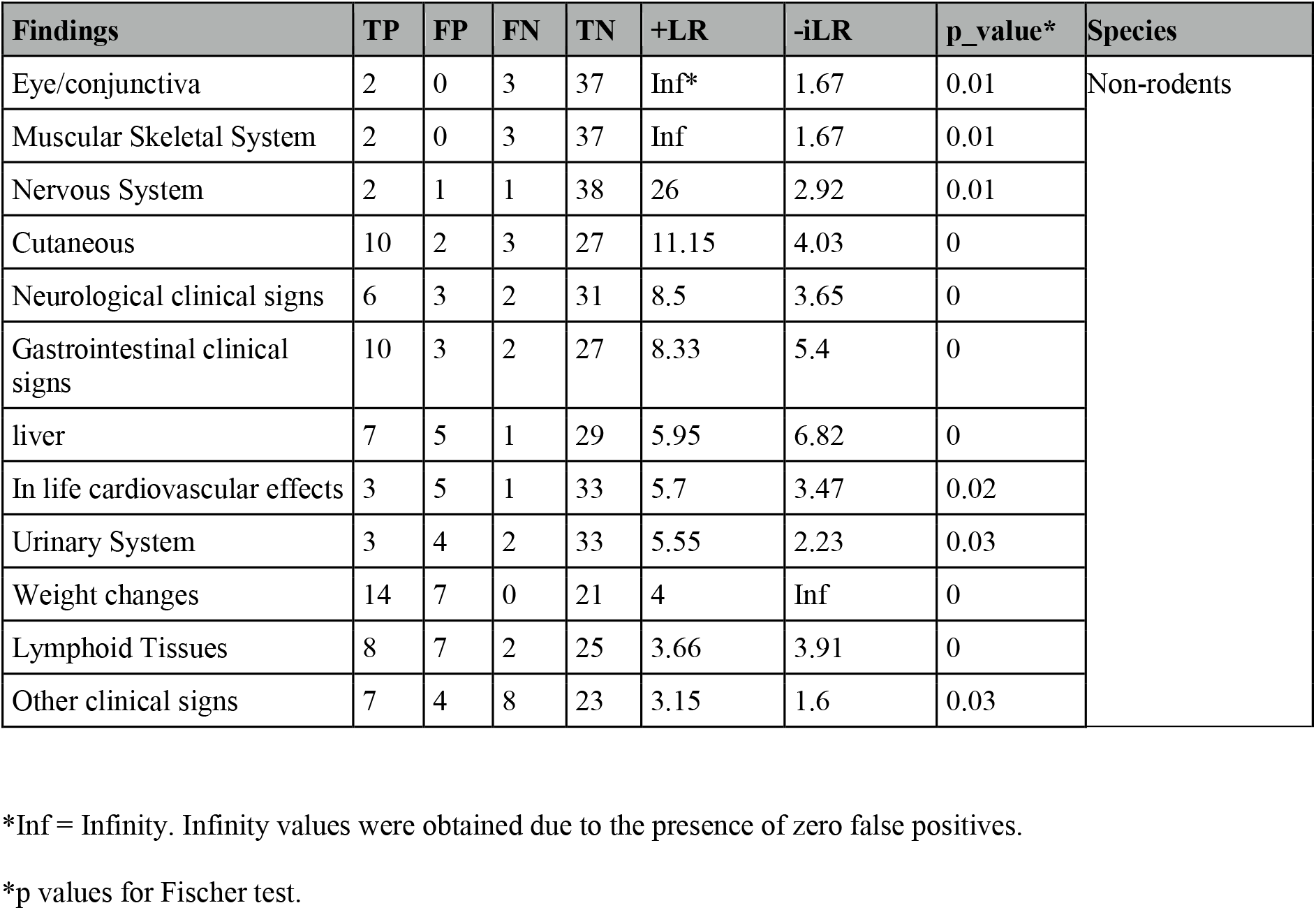
Values of contingency tables along with the statistically significant positive and negative inverse likelihood ratios of the non-adverse findings in non-rodents.

| Findings | TP | FP | FN | TN | +LR | -iLR | p_value* | Species |
| --- | --- | --- | --- | --- | --- | --- | --- | --- |
| Eye/conjunctiva | 2 | 0 | 3 | 37 | Inf* | 1.67 | 0.01 | Non-rodents |
| Muscular Skeletal System | 2 | 0 | 3 | 37 | Inf | 1.67 | 0.01 |  |
| Nervous System | 2 | 1 | 1 | 38 | 26 | 2.92 | 0.01 |  |
| Cutaneous | 10 | 2 | 3 | 27 | 11.15 | 4.03 | 0 |  |
| Neurological clinical signs | 6 | 3 | 2 | 31 | 8.5 | 3.65 | 0 |  |
| Gastrointestinal clinical signs | 10 | 3 | 2 | 27 | 8.33 | 5.4 | 0 |  |
| liver | 7 | 5 | 1 | 29 | 5.95 | 6.82 | 0 |  |
| In life cardiovascular effects | 3 | 5 | 1 | 33 | 5.7 | 3.47 | 0.02 |  |
| Urinary System | 3 | 4 | 2 | 33 | 5.55 | 2.23 | 0.03 |  |
| Weight changes | 14 | 7 | 0 | 21 | 4 | Inf | 0 |  |
| Lymphoid Tissues | 8 | 7 | 2 | 25 | 3.66 | 3.91 | 0 |  |
| Other clinical signs | 7 | 4 | 8 | 23 | 3.15 | 1.6 | 0.03 |  |

\*Inf = Infinity. Infinity values were obtained due to the presence of zero false positives.
\*p values for Fischer test.

**Table 7b.** Values of contingency tables along with the statistically significant positive and negative inverse likelihood ratios of the **adverse** findings in non-rodents.

| Adverse findings | TP | FP | FN | TN | +LR | -iLR | p_value | Species |
| --- | --- | --- | --- | --- | --- | --- | --- | --- |
| Nervous System | 2 | 0 | 0 | 40 | Inf | Inf | 0 | Non-rodents |
| Lymphoid Tissues | 2 | 1 | 0 | 39 | 40 | Inf | 0 |  |
| liver | 2 | 1 | 1 | 38 | 26 | 2.92 | 0.01 |  |
| Neurological clinical signs | 3 | 2 | 2 | 35 | 11.1 | 2.36 | 0.01 |  |
| Weight changes | 2 | 5 | 0 | 35 | 8 | Inf | 0.02 |  |

Less concordance was observed on the level of the -iLR of the non-rodent findings, where low values were observed for the Eye/conjunctiva, Muscular skeletal system, Nervous system, neurological clinical signs, In-life cardiovascular effects, urinary system, lymphoid tissues, and other clinical signs. A very high -iLR (equal to infinity) was observed for all the non-rodent adverse findings except for the liver and neurological clinical signs where low -iLR were observed (2.92 and 2.36 respectively) .

Lower +LR were observed for the majority of rodent findings (Table. 8a), except for the reproductive system, gastro-intestinal clinical signs, GIT and cutaneous findings. The same trend was observed for the -iLR. Regarding the adverse findings in rodents (Table. 8b) high +LR values were observed for all the adverse findings while lower -iLR values were observed for the liver and lymphoid tissues (-iLR < 5).

**Table 8a.** Values of contingency tables along with the statistically significant positive and negative inverse likelihood ratios of the non-adverse findings in rodents.

| Findings | TP | FP | FN | TN | +LR | -iLR | p_value | Species |
| --- | --- | --- | --- | --- | --- | --- | --- | --- |
| Reproductive System | 4 | 1 | 2 | 23 | 16 | 2.87 | 0 | Rodents |
| Gastrointestinal clinical signs | 7 | 2 | 2 | 19 | 8.17 | 4.07 | 0 |  |
| GI tract | 5 | 2 | 4 | 19 | 5.83 | 2.04 | 0.01 |  |
| Cutaneous | 5 | 3 | 3 | 19 | 4.58 | 2.3 | 0.02 |  |
| Endocrine System | 13 | 4 | 1 | 12 | 3.71 | 10.5 | 0 |  |
| liver | 14 | 4 | 2 | 10 | 3.06 | 5.71 | 0 |  |
| Lymphoid Tissues | 9 | 5 | 4 | 12 | 2.35 | 2.29 | 0.04 |  |
| Weight changes | 14 | 4 | 4 | 8 | 2.33 | 3 | 0.02 |  |

**Table 8b.**
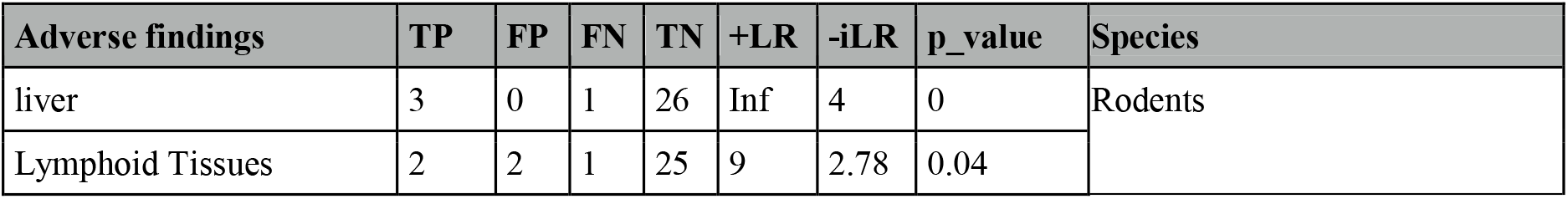
Values of contingency tables along with the statistically significant positive and negative inverse likelihood ratios of the **adverse** findings in rodents.

### Frequency of false positives and false negatives

Percentages of false positives and false negatives predicted by the short-term studies are represented for all the findings (Fig. 5) and adverse findings (Fig. 6). Similar to the likelihood ratios, only statistically significant contingency tables were analyzed and frequencies were calculated as explained in the Methods section. As seen in Fig. 4, frequencies of false positives and false negatives for findings in both rodents (Fig. 5A) and non-rodents (Fig. 5B) are all below 20%, denoting that the short-term studies successfully predicted the presence (True positives) or absence (True negatives) of the findings by 80%. Categories with false negative frequencies higher than 10% were: GI tract, lymphoid tissues and weight changes for rodent, while for non-rodents only the other clinical signs category showed a false negative frequency higher than 10%. Categories with false positive frequencies higher than 10% were weight changes, endocrine system, liver and lymphoid tissues for rodents and In-life cardiovascular effects, weight changes, liver and lymphoid tissues for non-rodents.

**Figure 5.**
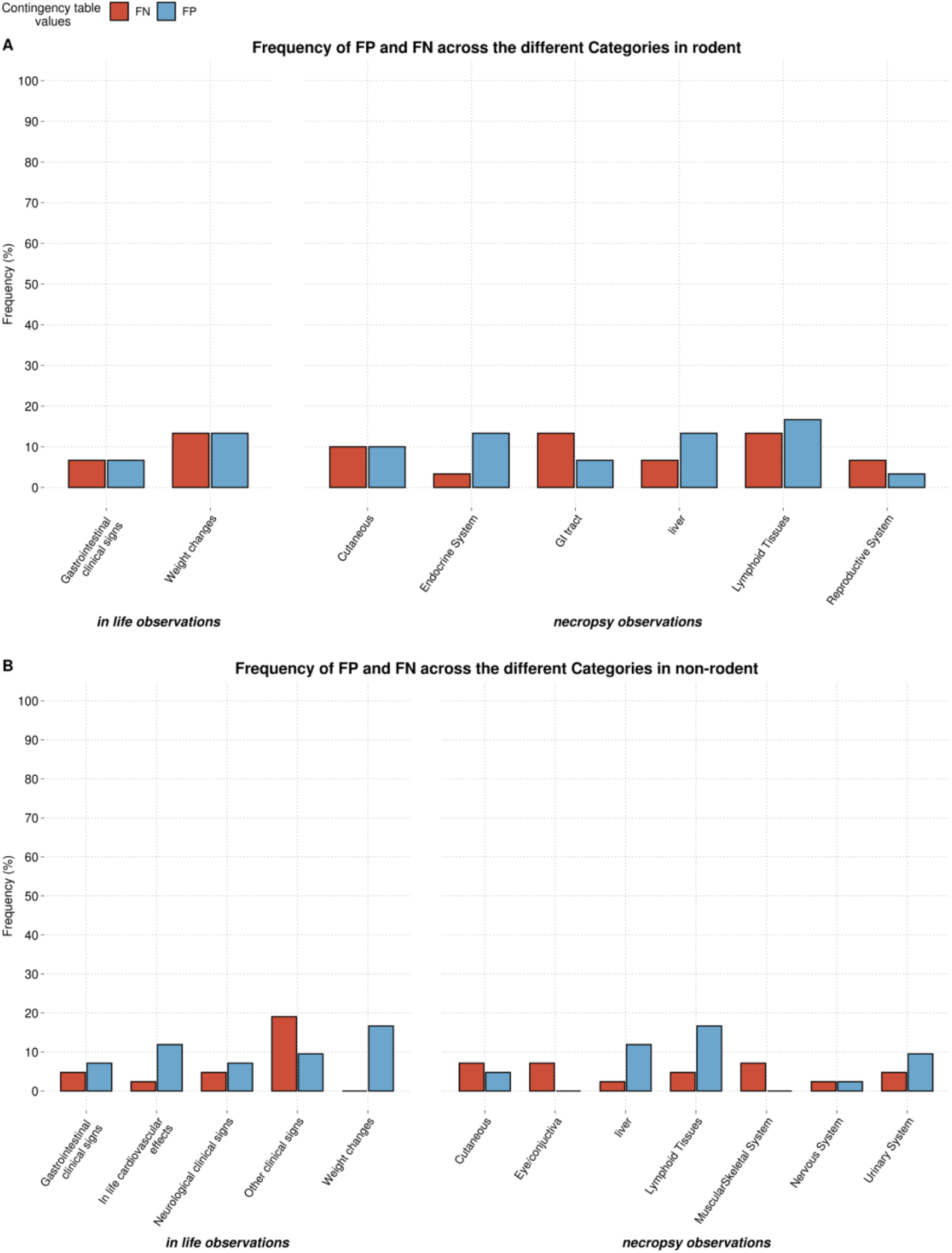
Frequency or percentage of false positives and false negatives of the findings across the different categories in (A) Rodents (B) Non-rodents

**Figure 6.**
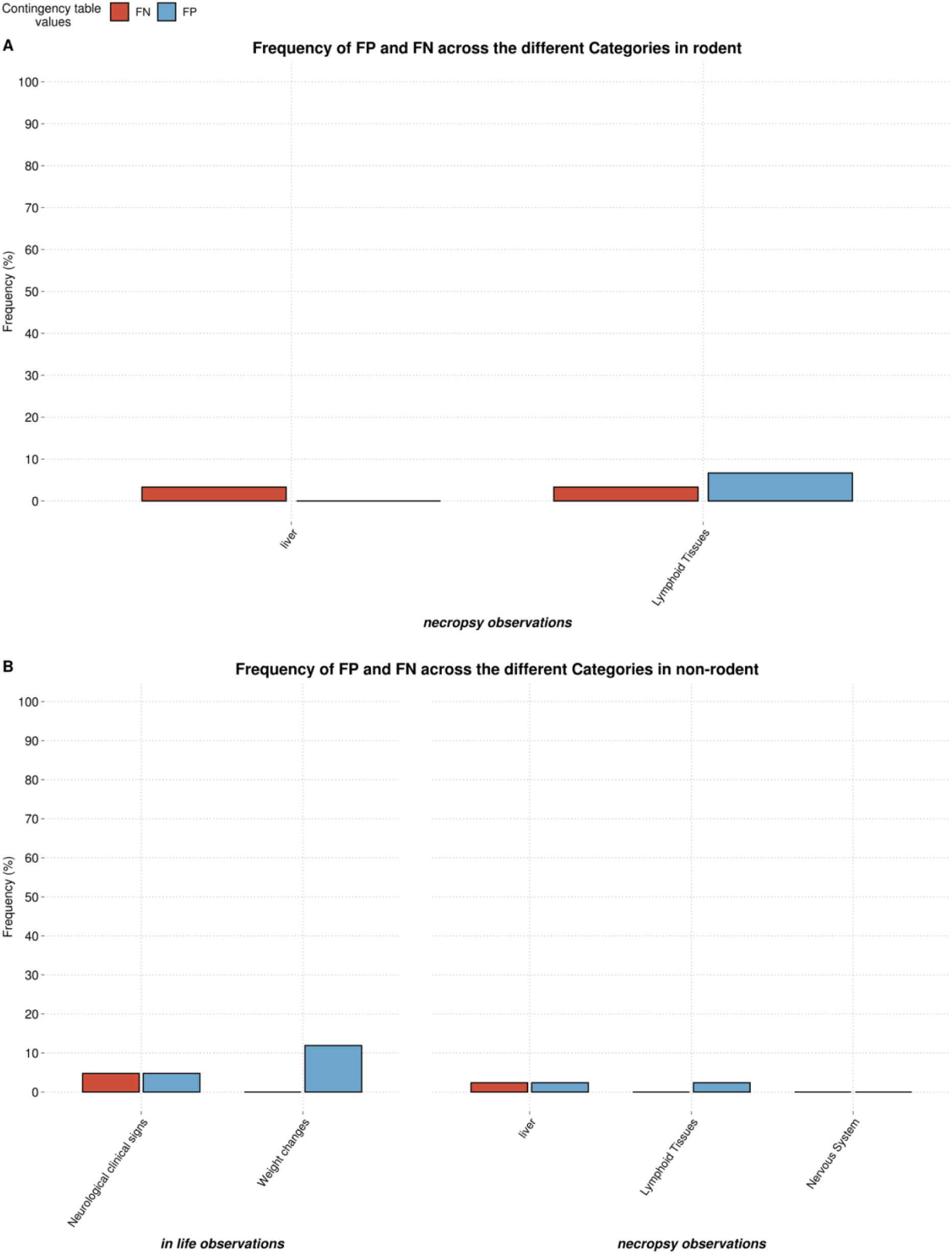
Frequency or percentage of false positives and false negatives of the **adverse** findings across the different categories in (A) Rodents (B) Non-rodents

In Fig. 6 frequencies of false positives and false negatives for the adverse findings are displayed for rodents (Fig. 6A) and non-rodents (Fig. 6B), all adverse findings showed frequencies less than 10% except for false positive weight changes in non-rodents.

Observations drawn from the false positive and false negative frequencies are in accordance with the likelihood ratios results. Where the overall results showed that the majority of the adverse findings exhibited high +LR and -iLR (except for lymphoid tissues adverse findings in rodent where -iLR = 2.78). However, it is of importance to analyze both measures, since they do not quantify miss-predictions similarly. The cut-off of “High” or “Low” likelihood ratios shows an impact on the results. For example, considering the adverse findings in rodents, low values of false positives and false negative frequency (6.6% and 3.3% respectively) were observed for the lymphoid tissue, indicating a good predictability of the short-term studies for the long-term adverse events. These frequencies might not be a sufficient indicator for the prediction capability of the short-term studies. Since upon investigation of the likelihood ratios, we observe that despite a high +LR being observed for both findings according to our chosen likelihood ratio cut-off, a low -iLR was (2.78) was seen. Indicating that there is a high probability to observe these adverse events in long-term studies, even if they were not observed by the short-term studies. A similar case can be observed upon the analysis of adverse findings in rodents, where liver and neurological clinical signs showed low false positive and false negative frequencies (2.3%) and (4.7%) respectively. While they exhibited low -iLR (2.92 and 2.36 respectively). Both false negatives and low -iLR are considered concerning in this analysis. Since the false negative indicates the presence of adverse events in the long-term studies and, their absence in the short-term studies. While a low -iLR indicates that if an adverse event was not seen in short-term studies, there is still a good probability that it will be seen in the long-term study.

## IV. Discussion

Minimization of the number of in-vivo studies is a challenging goal and requires the analysis of a vast amount of in-vivo data for high confidence conclusions to be made. Toxicity endpoints were extracted from 192 studies and a generally good concordance was seen between short-term and long-term studies on the overall adversity level and for most adverse findings (for small and large molecules). This number of studies however seemed to be insufficient to entirely address our main question, whether a study could be skipped. Nevertheless, the workflow adopted in the presented analysis was implemented as an open-source tool available for other researchers to use and adapt. In addition to that, we provide the dataset used in this analysis for reproducibility. This could facilitate the expansion of this analysis and its extrapolation to other datasets and, can therefore be considered as one of the main highlights of this study.

Another highlight of the study was the good predictivity for the adverse events between short and long-term studies in most of the cases, with only 1 out of 18 large molecules (Compound-O) and 2 out of 24 small molecules (Compound-6 and Compound-7) being unpredictive. Detailed analysis of the adverse events for these molecules showed concordance between the adverse findings of the short-term and mid-term studies for Compound-6, which gives rise to the question whether mid-term studies could be skipped. It might therefore be of importance to consider a concordance analysis between short-term and midterm findings in future studies. In addition to that, some of the unpredicted adverse events observed in the short-term studies were not considered adverse, but are known to increase in severity and incidence with longer study durations. Therefore, it might be of importance to consider “non-adverse” findings in assessing the predictability of short-term studies.

Secondly, analysis of the reasons behind the changes in the NOAEL was beneficial and might have a potential impact for decision making. For example, the NOAEL increase from short-term to long-term studies for the majority of the molecules was due to higher doses tested in the long-term studies. This suggests that adopting higher doses in short-term studies might provoke earlier manifestation of adverse events, minimizing the need for the long-term studies. Finally, analyzing both likelihood ratios and false positive/false negative frequencies was of importance due to the different significance given by each measure. It is also important to analyze the four quadrants of the contingency table and determine which measure fits best to the setting of the study, for example in our case it was important to analyze both the false negatives and false positives and not to limit the analysis to the true positives or true negatives Disregarding a false positive compound would result in terminating a compound for the wrong reasons, while disregarding a false negative would lead to missing adverse events risks upon skipping longer studies. Therefore, both measures constituted equal importance in this study. Multiple challenges were faced during this work, mainly being associated with the time-consuming nature of manual data extraction and its’ refinement. e in-vivo study reports are a valuable source of information on the safety findings of a drug candidate. Being frequently provided in PDF format, their manual extraction is required in addition to expertise in assessing the annotation of the findings as “treatment related” or “non-treatment related”. Attempting novel technologies for data extraction such as natural language processing and text mining techniques can enhance the cumbersome data extraction process, especially for the extraction of certain key findings such as study duration, dose, species and NOAEL [21, 29]. Standardization of the report formats is a prerequisite to facilitate the application of such techniques and obviate human intervention. Other similar challenges were encountered in this work, such as the verification of the controlled terminology of the findings, categorization of the effects and their attribution to the appropriate target organ system. A more systematic data extraction process would be needed for a large-scale application of the work.

Currently, the FDA requires a standard format for non-clinical data regulatory submission, known as “The Standard Exchange of Non-clinical Data” (SEND), which should facilitate the analysis of preclinical data in the future. Other efforts have been exerted in the standardization of data (with respect to pathology terms), such as the International Harmonization of Nomenclature and Diagnostic Criteria for Lesions in Rats and Mice (INHAND) [30]. Cross-institutional consortia and collaborations have also been exerted to address many of the previously mentioned challenges. For example, the eTOX project which highlighted the importance of legacy data sharing and leveraging the valuable in-vivo toxicity data existing in pdf formats within the archives of large pharmaceutical companies [31]. The e-TOX project was succeeded by the translational safety assessment consortium (eTRANSAFE), which covered many aspects of preclinical and clinical toxicity such as the translatability of in-vivo toxicities of a new chemical entity (NCE) into the clinical dimension and the extent or relevance of these findings to man, the application of text mining and other techniques for extraction, visualization and analysis of data from preclinical and clinical reports and prediction of safety events through computational tools [32]. Confidentiality and sensitivity of the in-vivo data is another challenge faced for large-scale application of such studies and was tackled by both the eTOX and eTRANSAFE projects.

Another important aspect in preclinical research is exploring the translation of adverse findings in animals to toxicities in human. Many studies have looked into the concordance between adverse findings in animal and first clinical trials, using preclinical studies as diagnostic tests for predicting outcome in human [24-26, 33, 34] and inconsistencies have been shown in the use of animal models in predicting human toxicities [35] .In this study our assessment of the ability of short-term studies to predict long-term effects of small and large molecules was limited to the comparison of findings from both durations. However, correlation and comparison of the short-term and long-term findings with the clinical adverse findings could also be a way, not only to minimize long-term studies, but to minimize in-vivo studies in general. Analysis of the clinical data corresponding to these molecules was not within the scope of this study and therefore should be considered in future work.

Many other works focused on the use of computer models for toxicity prediction and how to identify the knowledge gaps in this field [36–39]. Adopting innovative approaches for the prediction and extrapolation of time dependent findings like time series analysis models could also be considered as next steps for this work, however expanding the database and analyzing more studies using the developed workflow would be needed prior to the application of such predictive approaches.

## V. Conclusion

Minimization of the number of in-vivo studies can reduce animal use and can help to shorten the preclinical development timeline. In this work, we performed a statistical comparison between short-term and long-term toxicity endpoints and compiled the methodology applied in this work into an open-source software, available for other users. The software implements important functions that facilitate this comparison, such as an overview of the distribution of the effects exhibited in the studies, study durations and the species used in the analysis, calculation of the overall adversity of the molecules, the NOAEL changes of molecules and the likelihood ratios of the adverse findings. Although our analysis showed promising results, the major question remains unanswered due to the relatively small dataset size employed in this analysis. A larger number of molecules and expansion of the current dataset is required to reach a tangible conclusion that could potentially impact the toxicologists’ decisions on minimizing the study durations and number of studies performed. We encourage other researchers to implement, adapt and ameliorate this workflow and extend its application to public and in-house data.

## Supplementary material

**Table.S1** Detailed summary on all the findings extracted from the reports included under the high-level categories previously defined in Table 1

**Table.S2** Details on the microscopic pathological finding with respect to each target organ. These target organs constitute the high-level target systems explained in Table2.

**Table.S3** This table constitutes the main dataset used as an input to the workflow.

**Table.S4** This table constitutes the main pathology dataset used as an input to the workflow.

**Table.S5** Discrepancies in the NOAEL changes outcome between the algorithm and the toxicologist interpretation.

## Data and code availability

The CSL-Tox workflow is freely available for users and can be found at https://github.com/Roche/CSL-Tox. A detailed R markdown tutorial explaining the steps of the workflow is provided. The data necessary to reproduce this work is deposited in the same GitHub repository (**Table S3 and S4**).

## Authors contribution

All authors contributed to the study design. SD performed the data collection and curation. SD and DN implemented the framework. DN drafted the manuscript and analyzed the results based on all authors’ input. All authors read and approved the final version of the manuscript.

