## Supplementary material for "CSL-Tox: An open-source analytical framework for the comparison of short-term and long-term toxicity end points and exploring the opportunities for decreasing in-vivo studies conducted for drug development programs"

**Table S1.** Detailed summary on all the findings extracted from the reports included under the high-level categories previously defined in Table 1.

| Used Terminologies | Preferred Terminilogy |
|---|---|
| **(A) GIT clinical signs** | GIT clinical signs |
| vomitus | |
| emesis | |
| oral discharge | |
| fecal abnormalities | |
| reduced feces | |
| soft feces | |
| unformed feces | |
| white discoloration of feces | |
| diarrhea | |
| liquid feces | |
| excessive salivation | |
| salivation | |
| swollen abdomen | |
| bedding in mouth | |
| mouth rubbing | |
| **(B) Neurological clinical signs** | Tremors/convulsions |
| localised tremor | |
| tremor | |
| twitching | |
| increased seizure risk | |
| convulsive behavior | |
| clonic tonic convulsions with tremor | |
| convulsive episodes | |
| slow movements | Hyperactivity/hypoactivity |
| hypoactivity | |
| reduced activity | |
| decreased activity | |
| reduction in locomotor activity | |
| sedated behavior | |
| subdued behavior | |
| lethargy | |
| apathy | |
| reduced reaction to outside stimulus | |
| sedating effect | |

| | |
|---|---|
| unresposiveness | |
| increased activity | |
| agitation with atypical behavior | |
| retlessness | |
| rolling gait | abnormal gait/posture |
| stiff gait | |
| abnormal gait | |
| bent gait | |
| hunched gait | |
| impaired mobility | |
| abnormal mobility | |
| unsteady gait | |
| staggering gait | |
| staggered movement | |
| ataxia | |
| staggering ataxia | |
| limited usage of left hindlimb | |
| modified coordination | |
| incoordination | |
| paddling | |
| hunched posture | |
| abnormal posture | |
| stiff posture | |
| hunched back | |
| prominent backbone | |
| distorted body | |
| lying position | |
| recumbent posture | |
| lateral posture | |
| prone | |
| prone posture | |
| prostration | |
| recumbency | |
| tail elevation | |
| hunched position | |
| head movement | abnormal behavior |
| pushing their head through bedding | |
| side to side movement of the head | |
| yawning | |
| abnormal vocalisation | |
| whimpering | |

| | |
|---|---|
| vocalisation | |
| vocalising | |
| piloerection | piloerection |
| **(C)Other clinical signs** | |
| panting respiration | irregular respiration |
| irregular respiration | |
| deep breathing | |
| irregular breathing | |
| noisy breathing | |
| panting | |
| labored respiration | |
| discolored haircoat of the perioral region | abnormal haircoat |
| rough haircoat | |
| ruffled fur | |
| fur loss | |
| hairloss | |
| half closed eyes | partially closed or closed eyes |
| partially closed eyes | |
| semi closed eyes | |
| closed eyes | |
| poor condition | Morbidity |
| debilitation | |
| morbidity | |
| thin body | |
| thin appearance | |
| emaciation | |
| pale | |
| paleness | |
| moribund condition | |
| enterobacter cloacae infection in the skin | skin infection |
| skin thickening | skin thickening |
| flaky skin | skin lesions |
| skin lesions | |
| skin scales | |
| scabby skin | |
| focal skin lesions | |
| soiled areas | |

| | |
|---|---|
| multifocal skin lesions | |
| redenning of the extremities | |
| erythema of the ears and extremities | |
| wet lesion | |
| scabbing | |
| scabs | |
| red/black skin | |
| skin discoloration | Skin discoloration |
| black skin at eyelids | |
| black skin at periorbital areas | |
| swelling at the face | Face edema |
| swelling at eyelids | |
| swelling at periorbital areas | |
| swelling | |
| subcutaneous edema | |
| nasal discharge | discharge |
| eyes discharge | |
| lacrymation | |
| lacrymation and secretion | |
| cold extremities | hypothermia |
| cold to touch | |
| coldness to the touch | |
| decreased body surface temperature | |
| hypothermia | |
| decreased body temperature | |
| sunken eyeballs | dehydration |
| dehydration | |
| dehydration weakness | |
| depression of ERG amplitudes | ERG changes |
| ERG scotopic B wave amplitudes were depressed | |
| ocular inflammation | Ocular findings |
| perivascular sheathing | |
| hazy media | |
| vitreous cells | |
| limbal corneal pigmentation | |
| white particles in the aqueous humor | |
| brown particles in the aqueous humor | |
| white particles in the vitreous body | |
| reduced IOP | |

| | |
|---|---|
| increased retinal nerve fiber layer thickness | OCT changes |
| **(D)Vital Signs- Cardiovascular effects** | |
| increased heart rate | cardiovascular effects |
| decreased QT intervals | |
| RR interval decrease | |
| reduced heart rate | |
| bradycardia | |
| QTc prolongation | |
| increase in QTcV interval | |
| PR prolongation | |
| second degree atrioventricular block | |
| extrasystoles | |
| **(E)Macroscopic pathology** | |
| increase | increase |
| decrease | decrease |
| small | |
| firm | abnormal shape/surface |
| inflated | |
| raised area | |
| abnormal shape | |
| rounded | |
| granulated surface with medullary rays | |
| rough surface | |
| soft | |
| swollen | edema |
| edema | |
| cortical hemorrhage (adrenal gland) | hemorrhage |
| retinal hemorrhage | |
| subretinal hemorrhage | |
| hemorrhage | |
| pallor | discoloration |
| pale foci | |
| pale focus | |
| pale | |
| cortical pigmentation (adrenal gland) | |
| mottling | |
| mottled | |

| | |
|---|---|
| dark | |
| red discolourations | |
| red | |
| increase in grey and white and tan foc | |
| clay-like discoloration | |
| discolored | |
| discolored | |
| tan or white discoloration | |
| grey and white discoloration | |
| reddening of the glandular mucosa | |
| reddish colored foci in the glandular region | |
| thickened subcutaneous tissue and nodules | skin thickening, skin lesions |
| thickened subcutaneous tissue and scabs | |
| thickened subcutaneous tissue and nodules and hematomas and scabs | |
| thick sites | |
| sores | |
| lesions | |
| thickened | |
| skin thickening | |
| injection site mass | |
| scales | |
| ascites | effusions |
| hydropericardium | |
| hydrothorax | |
| distension | distension |
| calculus | calculus |
| erosions | erosions |
| decrease activity | decreased activity |
| increased adipose tissue (it refers again to neck, face, skin) | increased adipose tissue |
| prolonged persistence of injuries | prolonged persistence of injuries |
| dental abnormalities | dental abnormalities |
| tarsal hyperemia | tarsus bone hyperemia |
| vasculitis | vasculitis |
| decreased sensory conduction velocity | |
| decreased motor conduction velocity | |

**Table S2.** Details on the microscopic pathological finding with respect to each target organ. These target organ constitute the high level target systems explained in Table 2.

| Terminology | Controlled Terminology | Higher Level Grouping | |
|---|---|---|---|
| kidney | | | |
| tubular basophilic granules | tubular degeneration/regeneration | | |
| basophilic granules | tubular degeneration/regeneration | | |
| increased hyaline droplets | tubular hyaline droplets, increased | | brown seperate |
| degeneration/regeneration | tubular degeneration/regeneration | | |
| tubular foamy/granular macrophages | tubular foamy/granular macrophages | | |
| foamy macrophages | foamy/granular macrophages | | |
| basophilic granules in proximal tubular epithelial cells | tubular degeneration/regeneration | | |
| degeneration/regeneration of proximal tubular cells, | tubular degeneration/regeneration | inflammation/infiltrate | inflammation |
| pelvic inflammation | pelvic inflammation | | infiltrate |
| transitional cell hyperplasia in the renal pelvis | pelvic transitional cell hyperplasia | | |
| mononuclear cell infiltrate in the pelvic/peripelvic area | pelvic infiltrate | | |
| pyelonephritis | pyelonephritis | hypertrophy/hyperplasia | hyperkeratosis |
| pyelitis | pyelitis | | hypertrophy |
| tubular vacuolation | tubular vacuolation | | hyperplasia |
| increased levels of pigment | increased pigmentation | | |
| increased incidence of the chronic progressive nephropathy | chronic progressive nephropathy | | atrophy |
| extramedullary hematopoiesis | extramedullary hematopoiesis | | |
| tubular dilatation | tubular dilatation | | |
| epithelial attenuation and regeneration | epithelial attenuation, epithelial regeneration | | |
| inflammatory cell infiltration | inflammation, infiltrate | | congestion |
| proteinaceaous casts | Cast (tubular) | | hemorrhage |
| tubular basophilia | Basophilic tubule (tubular degeneration/regeneration) | | edema |
| nephropathy | chronic progressive nephropathy | | |
| tubular regeneration | Basophilic tubule (tubular degeneration/regeneration) | degeneration | degeneration/regeneration |
| degeneration and fibrosis | tubular degeneration/regeneration | | single cell necrosis/ apoptrosis |
| increased incidence of tubular basophilia | Basophilic tubule (tubular degeneration/regeneration) | | |
| increased incidence of hyaline droplets | tubular hyaline droplets, increased | adipocyte changes | adipocyte accumulation |

| | |
|---|---|
| tubular hyaline droplets | increased tubular hyaline droplets |
| glomerular vacuolation | glomerulopathy |
| glomerulopathy | glomerulopathy |
| adipocytes | adipocyte accumulation |
| urothelial lesions | pelvic lesions |
| hypertrophy of the lining epithelium of collecting ducts | epithelial degeneration/regeneration |
| tubular degeneration | tubular degeneration/regeneration |
| atrophy | atrophy |
| vacuolation | vacuolation |
| dilation | dilation |
| infiltrates of vacuolated macrophages | foamy/granular macrophages |
| intranuclear inclusions | intranuclear inclusions |
| glomerulonephritis | glomerulonephritis |
| tubular nephropathy | chronic progressive nephropathy |
| cortical atrophy | cortical atrophy |
| tubular atrophy with interstitial fibrosis | tubular atrophy, fibrosis |
| mixed cell infiltrates | infiltrate |
| granuloma formation | Granuloma |
| pelvic dilatation | pelvic dilatation |
| transitional cell hyperplasia | pelvic transitional cell hyperplasia |
| perirenal inflammation | perirenal inflammation |
| granular casts | cast (tubular) |
| inflammation | inflammation |
| granulomas | granuloma |
| vascular congestion | Congestion |
| congestion | Congestion |
| increase in the severity of hemosiderosis | Hemosiderin |
| **lymph node** | |
| basophilic granules | lymphoid degeneration/regeneration |
| increased sinus macrophages | macrophages |
| increased lymphoid cellularity | hyperplasia |
| vacuolated macrophages | foamy/granular macrophages |
| lymphoid depletion | lymphoid depletion |
| edema | edema |
| plasmocytosis | plasmocytosis |
| lymphocyte depletion | lymphoid depletion |
| necrosis | necrosis |
| lymphoid stimulation | lymphoid stimulation |

| | |
|---|---|
| Sinus histiocytosis | Sinus histiocytosis |
| **liver** | |
| granular kupfer cells | granular kupfer cells |
| hepatocellular single cell necrosis/apoptosis | hepatocyte apoptosis/single cell necrosis |
| hepatocyte basophilic granules | granular hepatocytes |
| basophilic granular kupffer cells | granular kupfer cells |
| kupffer cell hypertrophy | kupffer cell hypertrophy |
| hepatocyte hypertrophy | hepatocyte hypertrophy |
| increased mononuclear cell infiltrates | infiltrate |
| centrilobular hypertrophy | hepatocyte hypertrophy |
| centrilobular hepatocyte hypertrophy | hepatocyte hypertrophy |
| periportal hepatocellular vacuolation | hepatocyte vacuolation |
| increased mitoses | increased mitosis/increased mitotis figures |
| kupffer cell pigmentation | kupffer cell pigmentation |
| diffuse increase in lipid droplets | lipid accumulation |
| increased hepatocyte vacuolation | hepatocyte vacuolation |
| lipid accumulation | lipid accumulation |
| prussian blue positive pigment in kupffer cells | hemosiderin |
| increased incidence and severity of hepatocyte vacuolation | hepatocyte vacuolation |
| diffuse and centrilobular/midzonal hepatocellular lipid deposits | lipid accumulation |
| multinucleate hepatocytes | multinucleated hepatocytes |
| hepatocellular pigmentation | hepatocyte pigmentation |
| bile duct hyperplasia | bile duct hyperplasia |
| clear cell foci | foci |
| basophilic tigroid foci | foci |
| biliary hyperplasia | bile duct hyperplasia |
| portal inflammation | portal inflammation |
| yellow and brown pigments in kupffer cells | kupffer cell pigmentation |
| macrophages | macrophages |
| kupffer cell vacuolation | kupffer cell vacuolation |
| periacinar hypertrophy | hepatocyte hypertrophy |
| diffuse hypertropy | hypertrophy |
| periacinar vacuolation | hepatocyte vacuolation |
| periacinar lipid deposits | lipid accumulation |
| increase in the severity of hemosiderosis | hemosiderin |
| kupffer cell hemosiderin granules | Hemosiderin |
| diffuse hepatocellular lipid deposits, | lipid accumulation |
| centrilobular hepatocellular lipid deposits, | lipid accumulation |

| | |
|---|---|
| midzonal hepatocellular lipid deposits | lipid accumulation |
| shift of lipid deposits from periportal to periacinar or midzonal | lipid accumulation |
| increased frequency and severity of hepatocytic glycogen vacuolation | hepatocyte glycogen vacuolation |
| increased severity of pas staining for glycogen storage | hepatocyte glycogen vacuolation |
| hypocellularity of hematopoietic tissue | decreased hematopoiesis |
| increased hematopoiesis | increased hematopoiesis |
| basophilic granules in kupffer cells, | granular kupffer cells |
| eosinophilic intranuclear inclusions in hepatocytes | intranulcear inclusions |
| higher levels of glycogen vacuolation, | hepatocyte glycogen vacuolation |
| lower hepatic lipid levels, | lipid depletion |
| higher hepatocellular glycogen levels | hepatocyte glycogen accumulation |
| inflammatory cell foci | inflammation |
| glycogen depletion | hepatocyte glycogen depletion |
| decreased oil red-o fat staining | hepatocyte lipid depletion |
| multinucleated hepatocytes | multinucleated hepatocytes |
| tigroid basophilic foci | foci |
| cellular alteration | foci |
| hepatocellular adenoma | hepatocyte adenoma |
| increased extramedullary hematopoesis | extramedullary hematopoiesis |
| reduced glycogen content in hepatocytes | hepatocyte glycogen depletion |
| multifocal cytoplasmic granularity of hepatocytes | hepatocyte granularity |
| diffuse ito cell hyperplasia | ito cell hyperplasia |
| vascular congestion | congestion |
| extramedullary erythropoiesis | extramedullary hematopoiesis |
| hepatic glycogen vacuolation | hepatocyte glycogen vacuolation |
| kupffer cell hemosiderin granules | Hemosiderin |
| **Injection site** | |
| basophilic granules | basophilic granules |
| granular macrophages | foamy/granular macrophages |
| granulomatous inflammation | inflammation |
| subcutaneous fibrosis | fibrosis |
| collagen degradation | collagen degradation |
| subcutis necrosis | necrosis |
| edema | edema |
| hemorrhage | hemorrhage |
| mononuclear cell response | infiltrate |
| acute inflammation | inflammation |
| inflammatory cell infiltration | inflammation,infiltrate |

| | | | |
|---|---|---|---|
| increased perivascular inflammation | inflammation | | |
| collagen deposition | fibrosis | | |
| acanthosis | acanthosis | | |
| inflammatory lesions | inflammation | | |
| incidence and severity of inflammatory cell foci | inflammation | | |
| hyperkeratosis | hyperkeratosis | | |
| epidermal hyperplasia | epidermal hyperplasia | | |
| increased mononuclear, perivascular inflammatory cell foci | inflammation | | |
| inflammation | inflammation | | |
| subcutis inflammation | inflammation | | |
| subcutis adipose tissue atrophy | adipose tissue atrophy | Blood findings | hematopoiesis |
| mononuclear infiltrate | infiltrate | | |
| lymphoplasmacytic infiltrates | infiltrate | | |
| mixed infiltrate | infiltrate | | |
| hypertrophy of the muscle fiber in the skin and subcutis | hypertophy | | |
| perivascular infiltrate of mononuclear cells in the subcutis | infiltrate | | |
| perivascular infiltrate of mononuclear cells in the dermis | dermal infiltrate | | |
| serocellular crust (*2- would be nice to be clustered with something) | serocellular crust | | |
| **lung** | | | |
| equivocal low grade vascular changes | | | |
| increase in the incidence and grade of foamy macrophages | foamy/granular macrophages | | |
| increased infiltrate of intra-alveolar macrophages | macrophages | | |
| increased incidence of alveolar foamy macrophages | foamy/granular macrophages | | |
| congestion | congestion | | |
| **testis** | | | |
| tubular degeneration/atrophy | tubular degeneration/atrophy | | |
| granular macrophages | foamy/granular macrophages | | |
| increase of degenerated spermatocytes | spermatocyte degeneration | | |
| tubular degeneration | tubular degeneration/atrophy | | |
| degeneration of seminiferous tubules | tubular degeneration/atrophy | | |
| tubular vacuolation | tubular vacuolation | | |
| degeneration of germ cells | germ cell degeneration | | |
| multinucleate cells | multinucleate cells | | |

| | |
|---|---|
| **eye** | |
| retinal degeneration | retinal degeneration |
| multifocal degeneration of the retina | retinal degeneration |
| vacuolation of ganglion cell layer and inner nuclear layer and outer plexiform layer | retinal vacuolation |
| keratopathy | keratopathy |
| mixed cell infiltrate in the inner retina and optic disc and vitreous | infiltrate |
| perivascular plasma cell in the optic disc | infiltrate |
| mild-optic disc perivascular plasma cell | infiltrate |
| anterior uvea plasma cell | infiltrate |
| anterior uvea mononuclear infiltrate | infiltrate |
| limbus plasma cell | infiltrate |
| limbus mononuclear infiltrate | infiltrate |
| inner retina degeneration | retinal degeneration |
| inner retina pigmented macrophages | infiltrate? |
| **skin** | |
| epidermal hyperplasia | epidermal hyperplasia |
| epidermal single cell necrosis | epidermal single cell necrosis |
| inflammation | inflammation |
| erosion/ulcer | erosion/ulcer |
| serocellular crusts | serocellular crust |
| hyperkeratosis | hyperkeratosis |
| mixed inflammatory infiltrate | infiltrate, inflammation |
| apoptotic/dyskeratotic keratinocytes | abnormal keratinocytes |
| epidermal hyperplasia/hyperkeratosis | epidermal hyperplasia, epidermal hyperkeratosis |
| parakeratosis | abnormal keratinocytes |
| necrosis of the superficial layers of epidermis, | epidermal necrosis |
| epidermal and dermal inflammation | epidermal inflammation,dermal inflammation |
| focal, often ulcerative, dermatitis | ulcerative dermatitis |
| subcutaneous inflammation | subcutaneous inflammation |
| adipocyte accumulation | adipocyte accumulation |
| edema | edema |
| subcutis fat atrophy | subcutaneous adipose tissue atrophy |
| multifocal superficial ulcerative dermatitis | ulcerative dermatitis |
| subcutis-atrophy of the adipose tissue | subcutaneous adipose tissue atrophy |
| hypertrophy of the cutaneous muscle | cutaneous hypertrophy |
| epidermal necrosis | epidermal necrosis |
| dermal necrosis | dermal necrosis |
| single cell keratinocyte necrosis | single cell necrosis |

| | |
|---|---|
| **thymus gland** | |
| atrophy | atrophy |
| decreased lymphoid cellularity | atrophy |
| increased lymphoid cellularity | hyperplasia |
| decreased lymphocytes | atrophy |
| decreased cortical lymphocytes | atrophy |
| lymphoid atrophy | atrophy |
| increased lymphocytolysis | increased lymphocytolysis |
| delayed involution | delayed involution |
| diffuse atrophy | atrophy |
| lymphocyte depletion | lymphoid depletion |
| necrosis | necrosis |
| vascular congestion | congestion |
| thymic involution | involution |
| **pancreas** | |
| single cell necrosis | single cell necrosis |
| enlarged adipocytes | enlarged adipocytes |
| adipocyte accumulation | adipocyte accumulation |
| zymogen depletion | zymogen depletion |
| increased occurrence of apoptotic acinar cells | exocrine cell apoptosis |
| exocrine epithelial single cell necrosis | exocrine epithelial single cell necrosis |
| **colon** | |
| distension | distension |
| single cell necrosis | single cell necrosis |
| **epididymis** | |
| decreased sperm | decreased sperm |
| ductular epithelium degeneration/necrosis | degeneration/necrosis |
| **female reproductive system+male reproductive system** | |
| estrous cycle arrest | estrous cycle arrest |
| atrophy | atrophy |
| **Gastrointestinal tract** | |
| crypt epithelial single cell necrosis | mucosa epithelial apoptosis/single cell necrosis |
| degeneration/necrosis | degeneration/necrosis |
| apoptosis/single cell necrosis | apoptosis/single cell necrosis |

| | |
|---|---|
| dilation | dilation |
| mucosa edema | mucosa edema |
| submucosa edema | submucosa edema |
| crypt apoptosis | mucosa apoptosis |
| **bone marrow** | |
| decreased cellularity | hypocellularity |
| increase micronucleated erythrocytes | increased hematopoiesis |
| increased lymphoid cellularity | hyperplasia |
| lower incidence of fat in the sternum | adipose tissue atrophy |
| lower incidence of fat in the femur | adipose tissue atrophy |
| hypercellularity | hyperplasia |
| hyperplasia in the sternum | hyperplasia |
| fat reduction | adipose tissue atrophy |
| intracellular and extracellular perl's stained iron | hemosiderin |
| myeloid hyperplasia | hyperplasia |
| adipocyte accumulation | adipocyte accumulation |
| hypocellularity of hematopoietic tissue | hypocellularity |
| enlarged adipocytes | enlarged adipocytes |
| absence of erythroid precursors in sternum | decreased hematopoiesis |
| atrophy | hypocellularity |
| atrophy in the femoral adipose tissue | adipose tissue atrophy |
| increased hematopoiesis | increased hematopoiesis |
| increased production of nucleated erythrocytes | increased hematopoiesis |
| **tooth** | |
| odontoblast degeneration/necrosis | degeneration/necrosis |
| dental dysplasia | dysplasia |
| **hard palate** | |
| **epithelial single cell necrosis** | epithelial single cell necrosis |
| **mammary gland** | |
| atrophy | atrophy |
| increased intracellular brown granular pigment | increased pigmentation, granularity |
| diffuse lobuloalveolar hyperplasia | hyperplasia |
| acinar hyperplasia | hyperplasia |
| enlarged adipocytes | enlarged adipocytes |
| adipocyte accumulation | adipocyte accumulation |

| | |
|---|---|
| **thyroid gland (+ parathyroid gland)** | |
| follicular cell hypertrophy | follicular hypertrophy |
| follicular cell hyperplasia | follicular hyperplasia |
| diffuse follicular cell hypertrophy | follicular hypertrophy |
| follicular distension | follicular distension |
| increased incidence and severity of flattened thyroid gland follicular cell epithelium | follicular dilatation |
| adipocyte accumulation | adipocyte accumulation |
| follicular dilatation | follicular dilatation |
| follicular cell adenoma | follicular adenoma |
| **lymphoid tissues** | |
| increased lymphoid cellularity | hyperplasia |
| decreased lymphocytes | lymphoid depletion |
| **pituitary gland** | |
| mononuclear cell infiltrates | infiltrate |
| hypertrophy of cells of the adenohypophysis | hypertrophy |
| increased vacuolation | vacuolation |
| increased incidence of vacuolation | vacuolation |
| hypertrophy | hypertrophy |
| increased incidence of hypertrophy | hypertrophy |
| pars distalis hypertrophy | pars distalis hypertrophy |
| single cell hypertrophy | hypertrophy |
| **sciatic nerve+ nerve,tibial+nerve,trigeminal +ganglion,dorsal root+ hypoglossal nerve+glossopharyngeal nerve** | |
| mononuclear cell inflammation | infiltrate,inflammation |
| axonal degeneration | axonal degeneration |
| perineurium degeneration | perineurium degeneration |
| mononuclear infiltrate | infiltrate |
| axonal degeneration in the dorsal funiculus | axonal degeneration |
| vacuolation of neurons | neuron vacuolation |
| choroid plexus infiltrate | infiltrate |
| **urinary system** | |
| mononuclear cell infiltration | infiltrate |
| **spleen** | |
| increased lymphoid cellularity | hyperplasia |
| decreased lymphocytes | lymphoid depletion |

| | |
|---|---|
| increase in haemopoiesis | increased hematopoiesis |
| decreased germinal center and marginal zone cellularity | lymphoid depletion |
| vacuolated macrophages | foamy/granular macrophages |
| extramedullary hematopoiesis | extramedullary hematopoiesis |
| hemosiderosis | hemosiderin |
| increased pigment (iron) | hemosiderin |
| increased compensatory hemopoiesis | extramedullary hematopoiesis |
| increased hematopoiesis | increased hematopoiesis |
| infiltrates of vacuolated macrophages in the red pulp | foamy/granular macrophages |
| lower incidence of hematopoiesis | decreased hematopoiesis |
| decreased hematopoiesis | decreased hematopoiesis |
| necrosis | necrosis |
| vascular congestion | congestion |
| increased production of nucleated erythrocytes | increased hematopoiesis |
| congestion | congestion |
| **tonsil** | |
| lymphoid depletion | lymphoid depletion |
| increased lymphoid cellularity | hyperplasia |
| **larynx** | |
| epithelial hyperplasia | epithelial hyperplasia |
| parakeratosis | abnormal keratinocytes |
| apoptotic keratinocytes | abnormal keratinocytes |
| inflammation | inflammation |
| increased mitosis | increased mitosis (or mitotic figures) |
| apoptosis | apoptosis/single cell necrosis |
| single cell necrosis | apoptosis/single cell necrosis |
| ulceration and necrosis of the vocal fold epithelium | epithelial ulcer, epithelial necrosis |
| reactive squamous hyperplasia | epithelial hyperplasia |
| respiratory epithelium regeneration | epithelial regeneration |
| epithelial degeneration | epithelial degeneration |
| **ovary** | |
| increased corpora lutea | increased corpora lutea |
| increased vacuolation | vacuolation |
| adipocyte accumulation | adipocyte accumulation |
| **uterus** | |

| | |
|---|---|
| vacuolation | vacuolation |
| vacuolated cells | vacuolation |
| **jejunum+ileum+esophagus+conjuctiva** | |
| increased apoptosis, single cell necrosis | apoptosis/single cell necrosis |
| distension | distension |
| increased mitosis | increased mitosis (or mitotic figures) |
| inflammation | inflammation |
| single cell necrosis (mucosal epithelium) | mucosa epithelial single cell necrosis |
| mixed infiltrates | infiltrate |
| **adrenal gland** | |
| increased cortical microvacuolation (zona fasciculata but occasionally in the zona glomerulosa) | cortical vacuolation |
| lipid depletion | lipid depletion |
| increased cytoplasmic eosinophilia | lipid depletion |
| diffuse cortical hypertrophy | cortical hypertrophy |
| diffuse cortical vacuolation | cortical vacuolation |
| Diffuse hypertrophy of cortical zona reticularis | cortical hypertrophy |
| cortical atrophy | cortical atrophy |
| cortical hypertrophy | cortical hypertrophy |
| hypertrophy of the zona glomerulosa | cortical hypertrophy |
| cortical foamy alveolar macrophages | cortical foamy/granular macrophages |
| cortical vacuolation | cortical vacuolation |
| decreased vacuolation/cellular eosinophilia of the zona fasciculata | cortical vacuolation |
| adipocyte accumulation | adipocyte accumulation |
| decreased zona fasciculata | cortical atrophy |
| congestion | congestion |
| **brain** | |
| focal perivascular inflammation adjacent to the lateral ventricle | inflammation |
| extramedullary hematopoiesis | extramedullary hematopoiesis |
| lymphoplasmacytic infiltration of the choroid plexus | infiltrate |
| vascular congestion | congestion |
| vacuolation | vacuolation |
| epithelial vacuolation | epithelial vacuolation |
| **muscle,skeletal** | |

| | |
|---|---|
| degeneration/necrosis | myodegeneration/myonecrosis |
| hind limb myodegeneration/regeneration | myodegeneration/myoregeneration |
| myodegeneration | myodegeneration |
| myopathy | myopathy |
| myositis | inflammation |
| hypertrophy of the muscle fiber | hypertrophy |
| **heart** | |
| increased incidence and severity of the degenerative cardiomyopathy | cardiomyopathy |
| extramedullary hematopoiesis | extramedullary hematopoiesis |
| myodegeneration | myodegeneration |
| myonecrosis | myonecrosis |
| adipocyte accumulation | adipocyte accumulation |
| interstitial expansion | |
| cardiomyocyte hypertrophy | hypertrophy |
| distension | distension |
| myocardial hypertrophy | hypertrophy |
| brown adipose tissue changes | brown adipose tissue changes |
| white adipose tissue changes | white adipose tissue changes |
| cardiomyopathy | cardiomyopathy |
| mixed cell inflammation in the aorta root | inflammation, infiltrate |
| vascular congestion | congestion |
| hypertrophy of the muscle fiber | hypertrophy |
| **gland, harderian+gland,lacrimal** | |
| degeneration/atrophy | degeneration/atrophy |
| increased epithelial degeneration/regeneration | epithelial degeneration/regeneration |
| **parotid gland+salivary gland+salivary gland,sublingual+ submandibular gland+gland,brunner's** | |
| degeneration | degeneration |
| necrosis | necrosis |
| atrophy | atrophy |
| adipocyte accumulation | adipocyte accumulation |
| enlarged adipocytes | enlarged adipocytes |
| increase in incidence and severity of acinar vacuolation of the lingual salivary gland | vacuolation |
| increased secretory content | increased secretory content |
| mucin depletion in the acinar cells | atrophy |
| incomplete maturation | incomplete maturation |
| vacuolated macrophage infiltrate | foamy/granular macrophages |

| | |
|---|---|
| **Heart** | |
| aorta | |
| Epicardium | |
| macrovacuolation in brown fat | brown adipose tissue vacuolation |
| adipose tissue atrophy | adipose tissue atrophy |
| **mesentery** | |
| adipose tissue atrophy | adipose tissue atrophy |
| white adipose tissue diffuse atrophy | white adipose tissue atrophy |
| **stomach** | |
| mucosal/submucosal ulcers | erosion/ulceration |
| inflammation | inflammation |
| epithelial hyperplasia | epithelial hyperplasia |
| hyperkeratosis | hyperkeratosis |
| edema of the submucosa | submucosa edema |
| enlarged adipocytes | enlarged adipocytes |
| adipocyte accumulation | adipocyte accumulation |
| mucosa atrophy | mucosa atrophy |
| hyperkeratosis | hyperkeratosis |
| vascular congestion | congestion |
| glandular erosions | erosion/ulceration |
| mucosa erosions | mucosa erosions |
| mucosa hemorrhage | mucosa hemorrhage |
| lamina propria ulcer | erosion/ulceration |
| lamina propria necrosis | necrosis |
| **prostate gland** | |
| incomplete maturation | incomplete maturation |
| vacuolated macrophage infiltrate | foamy/granular macrophages |
| apoptosis in the urothelium of the urethra | apoptosis/degeneration urothelium/epithelium?? |
| **retroperitoneum+axilla+shoulder** | |
| univacuolar fat | white adipose tissue |
| brown adipose tissue changes | brown adipose tissue changes |
| increased lymphoid cellularity | hyperplasia |
| perirenal univacuolar fat | perirenal white adipose tissue |
| **bone+sternum** | |
| hyperostosis | hyperostosis |

| | |
|---|---|
| intramedullary formation | |
| hypocellularity | hypocellularity |
| hemorrhage | haemorrhage |
| sinus hyperemia | sinus hyperemia |
| hyperostosis | hyperostosis |
| atrophy | atrophy |
| haemorrhage | haemorrhage |
| **bladder+gallbladder** | |
| inflammation | inflammation |
| mixed inflammatory infiltrate | infiltrate |
| vacuolation | vacuolation |
| apoptosis | apoptosis/single cell necrosis |
| urothelium apoptosis, | urothelium apoptosis |
| urothelium vacuolation, | urothelium vacuolation |
| vacuolated macrophage infiltrate | foamy/granular macrophages |
| distension | distension |
| cholecystitis | inflammation (gall bladder though) |
| mixed infiltrate | infiltrate |
| urothelial hyperplasia, | urothelial hyperplasia (or epitheial hyperplasia, bladder) |
| uroliths | uroliths |
| **vagina** | |
| increased mucification | increased mucification |
| **oral cavity+tongue** | |
| ulceration in the mucosa | mucosa erosion/ulceration |
| single cell necrosis (mucosal epithelium) | mucosa epithelial single cell necrosis |
| mixed infiltrates | infiltrate |
| **Adipose Tissue, Brown** | |
| macrovacuolar (unilocular) changes, | vacuolation |
| increased and enlarged adipocytes | adipocyte accumulation, enlarged adipocytes |
| **ureter** | |
| dilatation | dilatation |
| mixed cell infiltrates | infiltrate |
| **nasal turbinate** | |
| brown pigment in the olfactory epithelium | pigment (epithelium) |

| | |
|---|---|
| **mucosa** | |
| lamina propria ulcer | lamina propria ulcer |
| lamina propria necrosis | lamina propria necrosis |

**Table.S3.txt** and **S4.txt** are provided as separate files in the GitHub repository https://github.com/Roche/CSL-Tox due to the size of the datasets.

**Table.S5** Discrepancies in the NOAEL changes outcome between the algorithm and the toxicologist interpretation.

| | **Species** | **Compounds** | **Algorithm** | **Toxicologist interpretation** |
|---|---|---|---|---|
| **Small molecules** | **Rodents** | Compound-9<br>Compound-25 | Decrease | Same |
| | **Non-Rodents** | Compound-8 | Increase | Same |
| | | Compound-21 | Increase | Decrease |
| **Large molecules** | **Rodents** | Compound-K | Increase | Same |
| | **Non-rodents** | Compound-K | Increase | Same |
| | | Compound-Q | Decrease | Same |
